# Individual repeatability, species differences, and the influence of socio-ecological factors on neophobia in 10 corvid species

**DOI:** 10.1101/2021.07.27.453788

**Authors:** Rachael Miller, Megan L. Lambert, Anna Frohnwieser, Katharina F. Brecht, Thomas Bugnyar, Isabelle Crampton, Elias Garcia-Pelegrin, Kristy Gould, Alison L. Greggor, Ei-Ichi Izawa, Debbie M. Kelly, Zhongqiu Li, Yunchao Luo, Linh B. Luong, Jorg J.M. Massen, Andreas Nieder, Stephan A. Reber, Martina Schiestl, Akiko Seguchi, Parisa Sepehri, Jeffrey R. Stevens, Alexander H. Taylor, Lin Wang, London M. Wolff, Yigui Zhang, Nicola S. Clayton

## Abstract

Behavioural responses to novelty, including fear and subsequent avoidance of novel stimuli, behaviours referred to as neophobia, determine how animals interact with their environment. Neophobia aids in navigating risk and impacts on adaptability and survival. There is variation within and between individuals and species, however, lack of large-scale, comparative studies critically limits investigation of the socio-ecological drivers of neophobia. In this study, we tested responses to novel objects and food (alongside familiar food) versus a baseline (familiar food alone) in 10 corvid species (241 subjects) across 10 labs worldwide. There were species differences in the latency to touch familiar food in the novel object and food conditions relative to the baseline. Three of seven socio-ecological factors influenced object neophobia: 1) use of urban habitat (vs not), 2) territorial pair vs family group sociality and 3) large vs small flock size (whereas range, caching, hunting live animals, and genus did not); while only flock size influenced food neophobia. We found that, overall, individuals were temporally and contextually repeatable (i.e. consistent) in their novelty responses in all conditions, indicating neophobia is a stable behavioural trait. With this study, we have established a network of corvid researchers, demonstrating potential for further collaboration to explore the evolution of cognition in corvids and other bird species. These novel findings enable us, for the first time in corvids, to identify the socio-ecological correlates of neophobia and grant insight into specific elements that drive higher neophobic responses in this avian family group.

## Introduction

Novelty is a common and vital aspect of animal life. The discovery of novel items and environments offers individuals an opportunity to benefit from new resources, such as food, tools, and shelter ^1,2^. Animals navigate novel stimuli through exploration, which allows for the assessment of any potential utility. However, novelty also presents the potential for danger: unknown food may be toxic, unknown objects may be traps and unfamiliar species may be predators ^1^. Consequently, various species also show fear and subsequent avoidance of novel stimuli, behaviours referred to as neophobia. Neophobia acts as a protective behaviour, encouraging hesitance and vigilance before/during exploration and thus helping to limit the danger associated with novelty ^1^. An appropriate level of neophobia within a species, according to their niche, should maximise their opportunity whilst minimising risk. As neophobia affects how animals interact with novelty, and novelty is a common occurrence, an understanding of neophobia is vital for animal cognition and behaviour research. This is particularly relevant as the world becomes heavily urbanised, with many species having to adapt to human-generated environmental changes and the inevitable novelty that follows ^3^. An understanding of the mechanisms underlying neophobia and any influencing factors may help explain why some species are more successful in adapting to new environments than others.

Previous research has investigated factors that may influence neophobia, as levels of neophobic behaviour vary between species and even individuals within a species (e.g. parrots ^4^ and ungulates ^5^). Many of these factors relate to socio-ecological factors, which may affect the costs and benefits of exploration and neophobia. However, there are very few large-scale comparative studies of neophobia, though one notable exception is Mettke-Hofmann et al. (2002) study on the relationship between a series of ecological factors, including diet and habitat, and both neophobia (latency to eat familiar food in presence of novel object) and exploration (latency to touch a novel object) behaviour in 61 species of parrot ^4^. The results suggested that a species’ ecology is closely associated with neophobia and exploration. Several different ecological variables influenced exploration, with species that inhabit complex habitats, have a diet of flower buds or fruits, and live on islands showing the shortest latencies in exploration tests. Two factors influenced neophobia: a diet of insects and a diet of leaves, indicating that parrots with a diet of insects were more neophobic than those feeding on plant material, explained as a possible consequence of the toxicity danger associated with insects ^4^. Thus, increased neophobia may mediate some of this risk. We note that this study did not test for individual repeatability over time or between conditions, used primarily small sample sizes (range 1-23 individuals, mean = 4.4, median = 2.5), and largely tested in uncontrolled social settings (e.g. measuring first individual to approach with/without others present) ^4^.

Many smaller-scale studies have investigated individual ecological factors that may affect neophobia within species. For example, individual common myna birds (*Acridotheres tristis*) who inhabit urban environments demonstrate lower levels of neophobia than those from rural areas and are quicker to utilise novel food resources ^6^. Greggor et al. (2016) found that wild birds (five corvid species, seven other bird species) approached human litter objects faster in an urban environment than in a rural environment ^7^. These findings have been suggested to occur because of habituation: birds in urban areas encounter human litter and objects more frequently than those in rural areas and thus become accustomed to this particular type of novelty. Other explanations have focussed on how urban areas offer low-risk and high-benefit environments, with a vast array of food resources in the form of human litter, and low levels of predation ^8^.

Differing habitats and diets may also influence neophobia and exploration. Greenberg and Mettke-Hofmann (2001) hypothesised that the costs of neophobia outweigh the benefits for generalist species, who utilise a range of resources that vary in availability, so reduced neophobia would allow for frequent exploration and discovery of new resources ^1^. However, specialist species, who use fewer, more stable resources, should show greater levels of neophobia as they have limited need to explore new food sources. This has been supported by research indicating that generalist Lesser-Antillean Bullfinch (*Loxigalla noctis*) showed shorter latencies to approach novel feeding stations than specialist bananaquit (*Coereba flaveola*) ^9^. Similarly, generalist song sparrows (*Melospiza melodia*) were less neophobic of objects than specialist swamp sparrows (*Melospiza georgiana*) in the field and in the lab ^10,11^.

Furthermore, social context, such as the presence of conspecifics, has been shown to reduce neophobia and increase exploration in several species. For example, zebra finches (*Taeniopygia guttata*) showed shorter latencies to eat from a novel feeder when in a flock than when alone ^12^. This may be due to group presence reducing generalised fear and/or risk being shared, thus reducing neophobia ^12^. It may also be context specific. For instance, Stöwe et al. (2006) found that ravens (*Corvus corax*) approached novel objects faster in the presence of siblings than non-siblings ^13^.

Ravens who are classed as “slow” explorers showed reduced latencies to approach novel objects when with a “fast” conspecific than when alone, but fast individuals’ approaches were impeded by conspecifics ^13^. Further, Chiarati et al. (2012) found that dominant breeding males in kin-based groups of carrion crows approached novel food before their other family members, reducing risks for their partner and offspring ^14^.

Individual differences in neophobia and exploration have been shown to be stable traits (i.e. repeatable or consistent over time and contexts) in some species, though inconsistent in others, which may be influenced by a range of factors, including the species, task, measures used, as well as seasonality, developmental, and social influences ^4,14–16^. Furthermore, although several socio-ecological variables appear to influence neophobia, a lack of large-scale comparative research limits interpretation of these effects (with the notable exception of ^4^), as well as testing whether it is a stable behavioural trait ^17^. Consistent methodology within a multi-species study allows for effective comparison within and between species ^18^, and thus would contribute towards understanding the mechanisms and influences of neophobia.

As a behavioural trait that dictates much of an animal’s interaction with the environment, including how they approach and solve novel problems, such data are valuable not only for establishing links between behaviour and ecology but also for studying cognition. Indeed, the time taken to learn a foraging task in feral pigeons (*Columba livia*) and zenaida doves (*Zenaida aurita*) covaried with individual levels of neophobia ^19,20^. Variation in neophobia also presents a potential confound for cognition research, as it can impact on performance in comparative cognitive tests, though is most often not tested or accounted for in relation to such comparisons between species ^20^. Outside of basic research, neophobia data may help inform applied animal welfare and conservation, including pre-release training in reintroduction programmes ^21^. For instance, working to increase neophobia levels in animals subjected to culling due to conflict with farmers ^21^.

Corvids (members of the crow family) are often featured in cognitive research ^22^, and are known to be relatively high on the scale of neophobia ^2,23^. Within corvids, species and individuals differ in neophobic propensities ^7,24–26^, as well as socio-ecological factors, such as range (how geographically widespread a species is), sociality, caching (hiding food for later use) behaviour, and tool-use ^22,27–31^. It is currently unknown what drives neophobia in corvids, for instance, whether they follow the same pattern as parrots relating to diet type e.g. ^4^, or whether there are different drivers of this variation. Corvids are therefore an optimal choice for these questions, however, to our knowledge, no study has yet compared neophobia comprehensively across many different corvid species, with repeated testing for individual repeatability, and directly testing the influence of socio-ecological factors.

We conducted a multi-lab collaborative study with three main aims: 1. compare neophobia across species 2. investigate the effect of socio-ecological factors on neophobia, and 3. assess individual temporal and contextual repeatability in neophobia. In 10 corvid species (241 subjects: Figure 1), we tested behavioural responses - specifically latency to touch familiar food – in three conditions: novel objects, novel food, and control condition (familiar food alone), with each condition repeated 3 times over 6-8 weeks (3 test rounds, 1 trial per condition per round, repeated every ∼2 weeks). Individuals were primarily tested while alone to control for any social influences and allow for repeated individual testing. Novel items were presented with familiar food to ensure behavioural responses were a result of the conflict between neophobia and desire for the familiar food, rather than, for example, exploration ^1^. Our response variable tested true food (and object) neophobia (i.e. fear of the appearance of the food), rather than dietary conservatism (i.e. latency to consume a novel food regularly in the diet) ^32^. We pooled resources across labs with the aim of increasing sample sizes and species representation. We selected tests that were not too time or labour intensive, given many labs were invited to contribute data, whilst giving a meaningful comparison across species that is largely based on established methodologies (i.e. latency to eat/ approach familiar food in the presence of a novel item).

**Figure 1.**
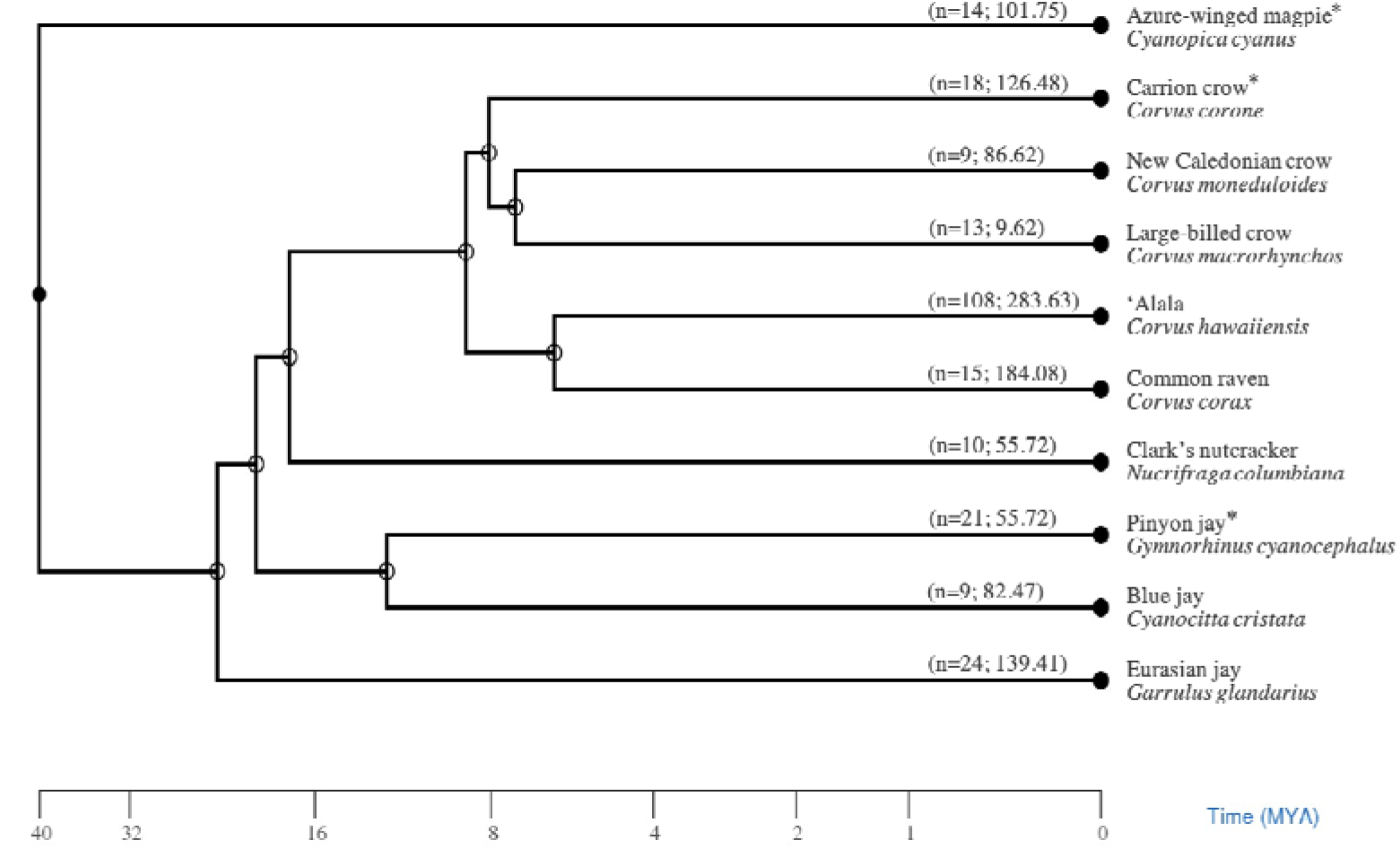
Phylogenetic tree. Sourced from http://www.timetree.org with sample size (n=x) and relative object neophobia score per species (mean latency to touch familiar food difference score i.e. novel object minus control value) - shigher score indicates higher neophobic response to novel object. * donates species tested at 2 sites

Firstly, we compared food and object neophobia between corvid species. We expected to find some species differences, as indicated by previous comparative corvid research e.g. ^7,25^. Next, we tested for the influence of socio-ecological factors: range (broad vs restricted/endemic), use of urban habitats (in addition to suburban/rural), hunting live animals, adult sociality (territorial vs family groups), flock size (small vs large), food caching (moderate vs specialised), and genus (*Corvus* or not) on neophobia. We expected that, like diet in parrots ^4^, neophobia would relate closely to aspects of species ecology. Specifically, in line with some previous research, we expected that species inhabiting a broad range, and utilising urban habitats, would show lower neophobia compared to those in restricted ranges and using only sub-urban/rural areas ^6–9,11,33^. Lower neophobia was also expected from species that live in larger flocks and family groups compared to small flocks and territorial pairs, due to the potential of risk-sharing between larger groups ^12,34^. As the influence of live hunting (selected as the species tested were otherwise generalists), caching and genus have not been previously tested in similar species, we had no a priori predictions for these factors. Finally, we tested for individual temporal and contextual repeatability. We expected to find individual repeatability, as there were only short delays between test rounds (∼2 weeks), similar to a related study in ‘A (*Corvus hawaiiensus*)^34^.

## Results

### 1. Species differences

Latency to touch familiar food differed across conditions (LMM: X^2^=316.05, df=2, p<0.001), test rounds (X^2^=28.75, df=1, p<0.001), and species (X^2^=93.03, df=9, p<0.001). The birds waited longer with a novel object or novel food present compared to the control condition (Tukey contrasts: novel object – control, z=18.79, p<0.001; novel food – control, z=7.97, p<0.001), and they waited longer when a novel object was present than when a novel food was present (z=7.35, p<0.001) (Figure 2). While latency to touch familiar food did not differ between rounds 1 and 2 (Tukey contrasts: z=0.57, p=0.371), it decreased in round 3 compared with round 1 and 2 (rounds 1 – 3, z=4.94, p<0.001; rounds 2 – 3, z=4.35, p<0.001) (S1 Figure). We also found that latency differed across species (S1 Table; Figure 3).

**Figure 2.**
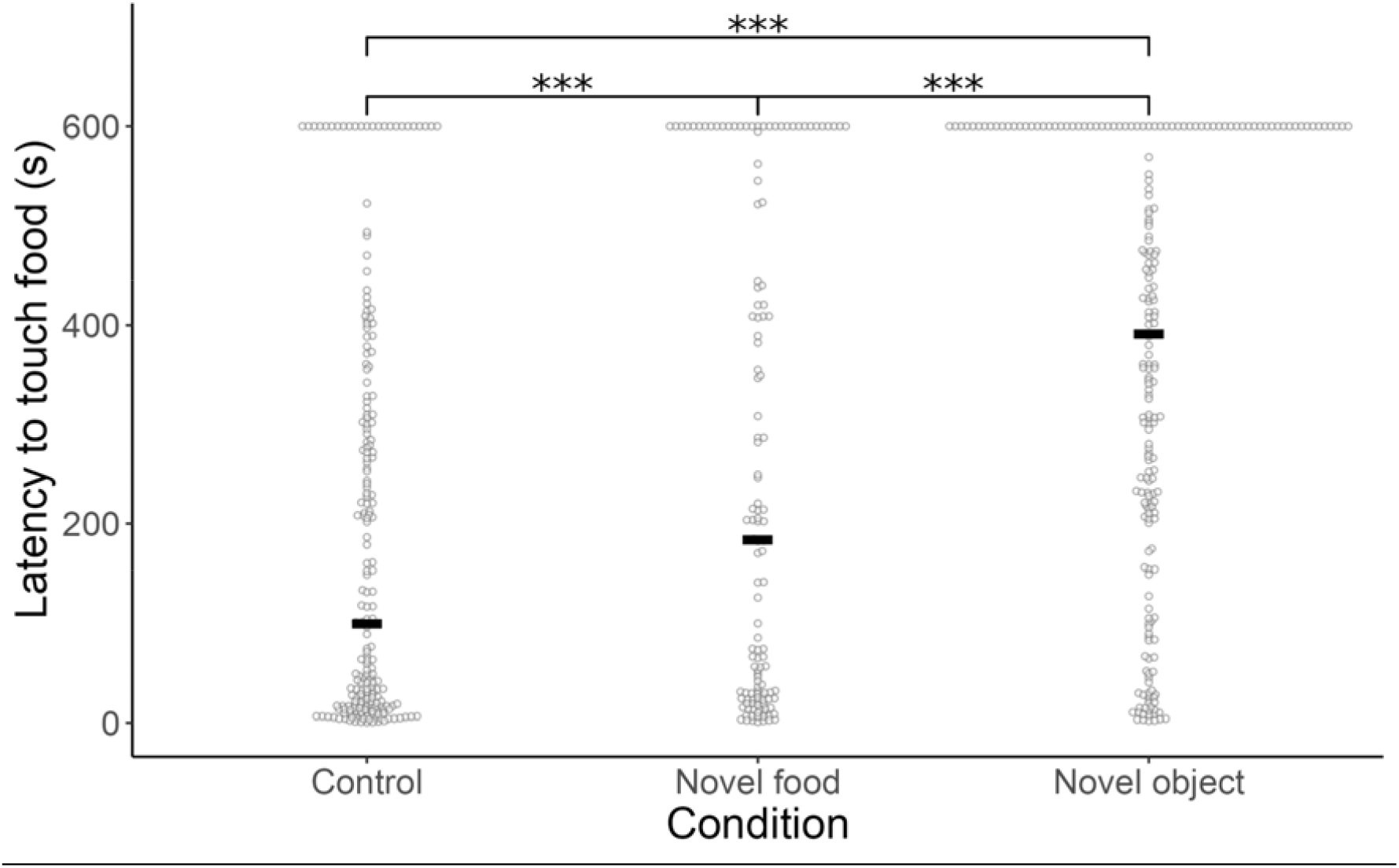
Latency to touch familiar food in each condition across all species. Control, novel food, and novel object conditions all differed from each other. Points represent individuals, lines represent median. *** p < 0.001

**Figure 3.**
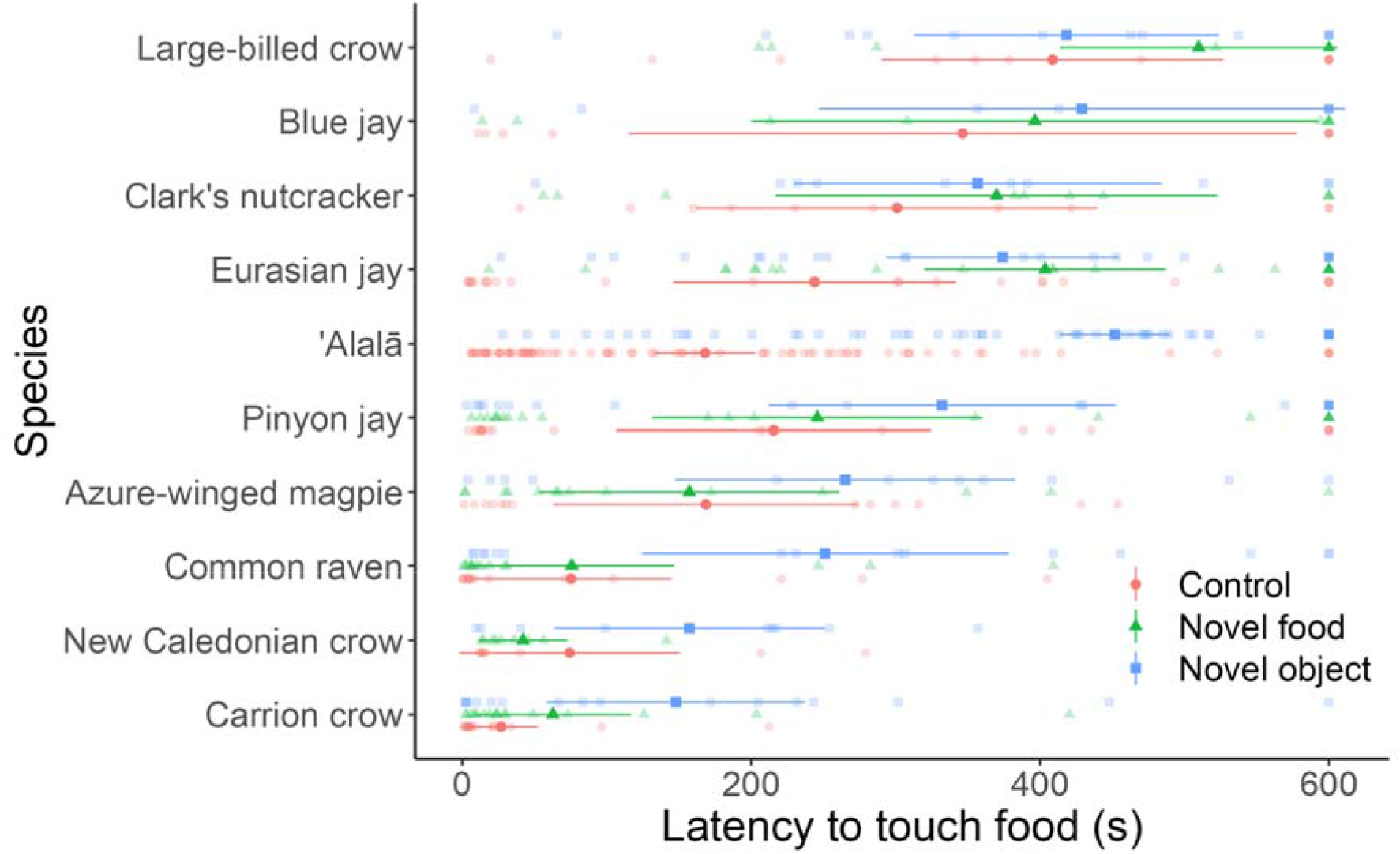
Latency to touch familiar food in each condition for each species. Some species differed in mean latency. Individual points represent subject means over rounds, points with error bars represent species means and 95% confidence intervals.

However, a potential confound of this study is that most species were housed and tested at different sites and therefore site is largely correlated with species. Three species were tested at two different sites. Using exploratory analysis, within these three species, we found that site did not affect latency to touch familiar food in carrion crows or azure-winged magpies but did affect latency in pinyon jays (S2 Table; S2 Figure).

To aid in standardizing latencies across sites as well as control for baseline neophobia and current motivational state, we created pairwise difference scores by subtracting the control latencies from the novel object and novel food latencies for each round and individual. Positive difference scores represent slower approaches to familiar food when a novel object/food is present (neophobia) and negative difference scores represent faster approaches (neophilia). The novel object difference scores differed across species (LMM: X^2^=47.02, df=9, p<0.001) and round (X^2^=8.18, df=1, p=0.017), with some differences between pairs of species (S3 Table; Figure 4A). Using novel object difference scores, common ravens were more neophobic than azure-winged magpies, large-billed crows, New Caledonian crows, Clark’s nutcrackers, blue jays and pinyon jays; azure-winged magpies, pinyon jays and Eurasian jays were more neophobic than large-billed crows; Eurasian jays were more neophobic than blue jays and Clark’s nutcrackers; carrion crows were more neophobic than Clark’s nutcrackers and large-billed crows; ‘Alala□ were more neophobic than blue jays, large-billed crows, Clark’s nutcrackers, New Caledonian crows, pinyon jays (Figure 4A).

**Figure 4.**
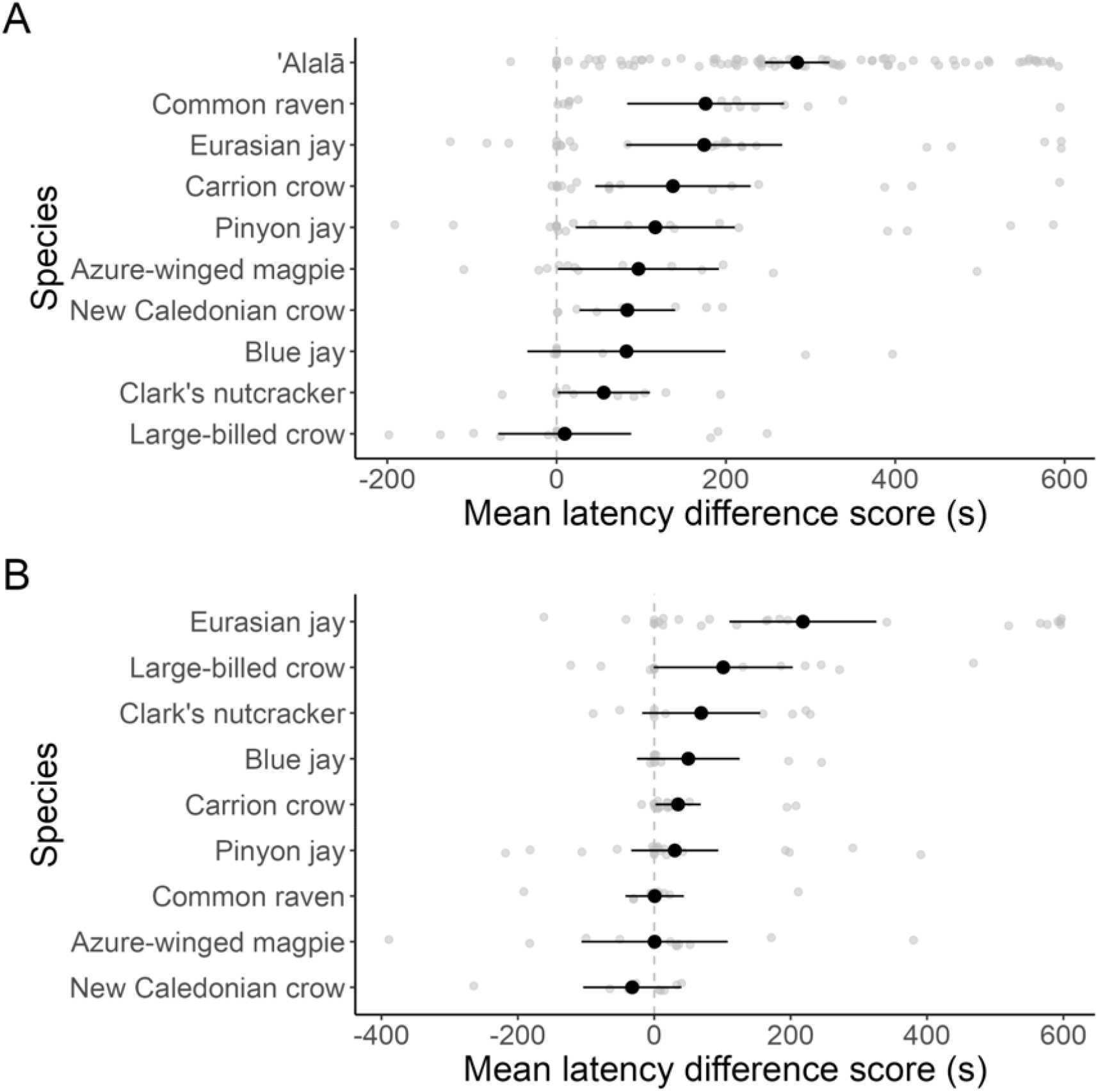
Species comparison using difference scores. Mean latency difference scores varied across species for (A) novel object neophobia and (B) novel food neophobia. Positive difference scores represent slower approaches to familiar food when a novel item was present (i.e. neophobia) and negative difference scores represent faster approaches (i.e. neophilia). Points represent individuals.

The novel food difference scores also differed across species (^2^=23.49, df=8, p=0.003) but not round (^2^=5.58, df=2, p=0.062). Note that ‘A s were not tested in the novel food condition and are removed from this analysis. Using novel food differences scores, Eurasian jays were more neophobic than all other species (Figure 4B; S4 Table). Overall, for both object and food conditions, most species were neophobic with mean difference scores greater than 0, with only New Caledonian crows showing a negative mean difference score for the food condition.

### 2. Effect of socio-ecological factors

Using novel object difference scores, object neophobic responses were affected by urban habitat use (X^2^=7.23, df=1, p=0.007), adult sociality (X^2^=6.61, df=1, p=0.010), and flock size (X^2^=4.98, df=1, p=0.026), but not range (X^2^=0.59, df=1, p=0.443), caching (X^2^=2.78, df=1, p=0.100), live hunting (X^2^=2.36, df=1, p=0.125), or genus (X^2^=0.24, df=1, p=0.628). Specifically, species that use urban habitats (as well as other habitats), live in larger flocks and family groups were less neophobic than those that do not/ very limited use of urban habitats, live primarily in territorial pairs or in smaller flocks (Figure 5A). Using novel food difference scores, food neophobia was only affected by flock size (X^2^=8.99, df=1, p=0.003) and not range (X^2^=2.72, df=1, p=0.100), urban habitat (X^2^=0.33, df=1, p=0.564), adult sociality (X^2^=1.98, df=1, p=0.160), caching (X^2^=0.25, df=1, p=0.621), live hunting (X^2^=0.10, df=1, p=0.756), or genus (X^2^=3.55, df=1, p=0.060). In contrast to the object neophobia finding, species that typically live in small flocks were less neophobic of novel food than those living in large flocks (Figure 5B).

**Figure 5.**
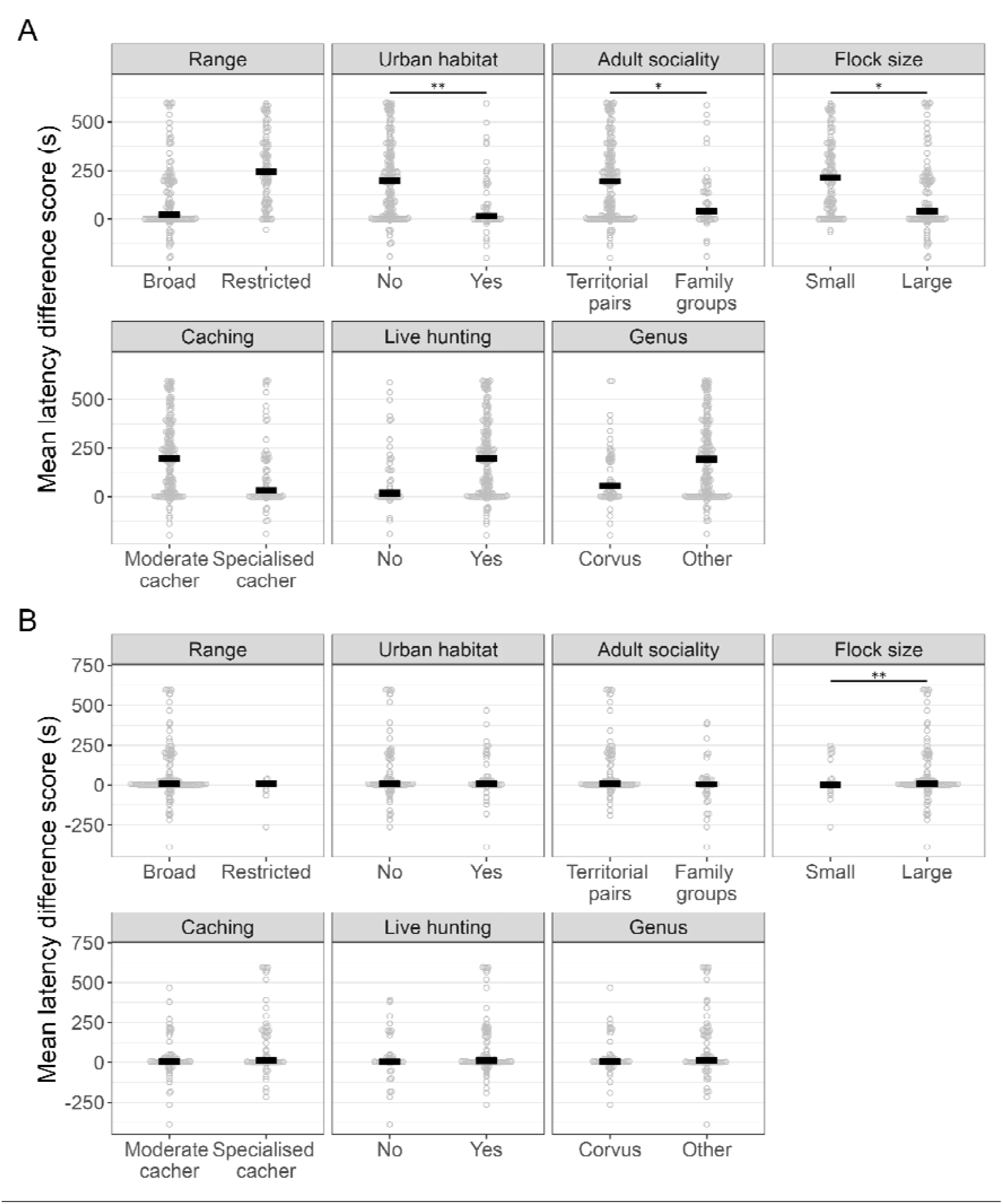
Effect of socio-ecological factors on neophobia. Linear mixed model on socio-ecological factors affecting latency to touch familiar food, using difference scores showed effects of urban habitat, adult sociality, and flock size on novel object neophobia (A) and effect of flock size on novel food neophobia (B). Positive difference scores represent slower approaches to familiar food when a novel object is present (i.e. neophobia) and negative difference scores represent faster approaches (i.e. neophilia). Points represent individual subjects and horizontal bars represent medians.

### 3. Individual temporal and contextual repeatability

Across all species, individuals’ responses to novel stimuli were temporally repeatable across test rounds (1-3) and contextually repeatable across conditions (control, novel object, novel food) (intra-class correlation coefficient: N = 217, ICC =0.462, p<0.001, CI = 0.402-0.521). In addition, responses were temporally repeatable within each condition (control: N = 216, ICC =0.542, p <0.001, CI = 0.467-0.625; novel object: N = 215, ICC =0.548, p <0.001, CI = 0.469-0.625; novel food: N = 132, ICC =0.477, p <0.001, CI = 0.381-0.591) (S5 Table). A within-species analysis showed similar temporal repeatability except for the New Caledonian crows (all conditions), azure-winged magpies (novel food only) and large-billed crows (novel object only), with contextual repeatability in all species except for the New Caledonian crows (S5 Table, S6 Table). Note that ‘Alala□ were not tested in the novel food condition.

## Discussion

In our multi-lab collaborative study, we tested the responses (latency to touch familiar food) of 10 corvid species to novel objects and food (beside familiar food), compared with a control baseline condition (familiar food alone). We found: a) some species differences in latency to touch familiar food in the presence of a novel object or novel food relative to baseline, b) effects of three socio-ecological factors - urban habitat use, adult sociality, and flock size - on object neophobia, and an effect of flock size on food neophobia, and c) individual temporal and contextual repeatability across species, as well as within species for all species except New Caledonian crows (all conditions), azure-winged magpie (novel food) and large-billed crow (novel object). The novel object and novel food conditions elicited higher neophobic responses (i.e. higher latencies) than the control condition. Additionally, neophobic responses reduced across rounds, with lower latencies in round 3 of testing than either round 1 or 2.

Species differed in object and food neophobia. For instance, we found that: common ravens, ‘Alalal□ and Eurasian jays were more neophobic than most other species tested for object neophobia, with Eurasian jays being more neophobic than all other species for food neophobia (using difference scores). The mean difference scores showed primarily neophobic responses to novel items (i.e. positive scores) compared to neophilic responses (i.e. negative scores). The critical test for interpreting these species differences, which is not possible in most of the previous research with single or small numbers of species/ individuals, was to test for specific influences of several socio-ecological factors that naturally differ between these corvid species. We found that three of seven factors tested influenced object neophobia: urban habitat use, adult sociality and flock size, while range, caching, hunting live animals and genus did not. Specifically, object neophobia was lower in species using urban habitats (n=5 species), living in family groups (n=3) and large flocks (n=6) compared with those only using suburban/ rural areas (n=5 species), living primarily in territorial pairs (n=7), or small flocks (n=4). Only flock size influenced food neophobia, with those living in small flocks showing lower neophobia than those in large flocks.

We expected urban habitat use to influence neophobia, based on previous research in other species, such as within-species comparisons in common myna ^6^ and black-capped chickadees ^33^. Urban habitats typically provide environments that are rich in novel stimuli, including human litter and manmade structures. Consequently, individuals and species inhabiting these areas are frequently exposed to various types of novel objects and may become habituated to such novelty. The costs of neophobia may also outweigh the benefits in urban habitats: human objects may become useful resources (litter may contain food or be an effective tool), an opportunity that would be lost by a high neophobic response. Additionally, urban environments have a relatively low predation risk for corvids and other animals, thus limiting the dangers associated with exploration of novel objects ^8,12^.

Similarly, we expected sociality to influence neophobia, with lower object neophobia in large flocks or family groups due to increased risk-sharing, compared with species living primarily as territorial pairs while adult or small flocks ^12^. Social presence has been shown in some species, including corvids, to have either a facilitating or inhibiting effect on neophobia and exploration ^15,27,35^. We differentiated species as ‘territorial’ vs ‘family groups’ according to their most prevalent social organisation ^28^. Some of these species do have quite flexible systems based on fission-fusion, such as common raven ^36^, thereby, they may be territorial as adults and/or during breeding season but be fairly tolerant of one another as juveniles or outside of breeding season ^37^. We therefore included a second sociality related factor: ‘small’ (up to 100 individuals) vs. ‘large’ flocks (over 100 individuals). It is interesting to note that we see contrasting effects of flock size on object compared with food neophobia, and effects of sociality even with individual testing (i.e. tested while alone).

We did not find an effect of hunting live animals on food or object neophobia (hunting live animals n=6 species vs not n = 4), which was our main dietary related measure, as otherwise, these corvids are largely similar in their diets. We may see a stronger effect of this factor with different types of novel food or in predator neophobia tasks. There was no effect of caching, despite differences between moderate (n=6 species) and specialised cachers (n=4) in the amount and type of food items that they cache. Our caching differentiation was based on a categorization of food caching into low, moderate, and specialized species ^38^ (Table 1), though it should be noted that some corvids also cache objects ^38,39^. However, there was insufficient prior data available to differentiate all species according to variation in the amount and type of object caching. Should this data become available in future, it would be worth testing our data to explore whether object cachers also differ in neophobia.

**Table 1.** Socio-ecological factors of corvid species tested.

| Species | Range | Urban habitat | Hunting live animals | Food caching | Adult sociality | Flock size |
| --- | --- | --- | --- | --- | --- | --- |
| Common raven, *Corvus corax* | Broad | *No | Yes | Moderate | Territorial pairs | Large |
| Carrion/ hooded crow, *Corvus corone*; *C. cornix* | Broad | Yes | Yes | Moderate | **<br>Territorial pairs | Large |
| Large-billed crow, *Corvus macrorhynchos* | Broad | Yes | Yes | Moderate | Territorial pairs | Large |
| New Caledonian crow, *Corvus moneduloides* | Restricted | No | No | Moderate | Family groups | Small |
| Alala□, *Corvus hawaiiensis* | Restricted | No | Yes | Moderate | Territorial pairs | Small |
| Eurasian jay, *Garrulus glandarius* | Broad | Yes | Yes | Specialised | Territorial pairs | Large |
| Pinyon jay, *Gymnorhinus cyanocephalus* | Broad | No | No | Specialised | Family groups | Large |
| Blue jay, *Cyanocitta cristata* | Broad | Yes | No | Specialised | Territorial pairs | Small |
| Clark's nutcracker, *Nucifraga columbiana* | Broad | No | Yes | Specialised | Territorial pairs | Small |
| Azure-winged<br>magpie,<br>*Cyanopica*<br>*cyanus* | Broad | Yes | No | Moderate | Family<br>groups | Large |

Differentiation within factors restricted to 2 levels reflecting availability of published data to support these distinctions across all species. \* Typically applicable for Europe (where the common ravens tested in this study were held and sourced); ravens have used/use cities at some North American sites (personal communication, Thomas Bugnyar). \*\* One carrion crow population in Spain have helpers at the next (i.e. cooperative breeding), though this is not reported in other populations<sup>48</sup>

We found no effect of range (broad n=8 vs restricted n=2 species) on either food or object neophobia, which was unexpected, according to the “island tameness theory”, which suggests that island populations may be less neophobic because they have evolved with fewer dangers in the environment ^40^. We note, however, that only the New Caledonian crows and ‘Alalal□ had a restricted i.e. endemic range, therefore interpretation of this finding should be tentative, particularly as the New Caledonian crows were wild sourced. Finally, we found no effect of genus (Corvus n=5 or not n=5 species) on neophobia. Should additional reliable phylogenetic data for corvids become available, and we were able to increase the number of species above 20 species ^16,41,42^, we should be able to include further phylogenetic controls in future.

All species, other than New Caledonian crows (all conditions), azure-winged magpies (novel food) and large-billed crows (novel object) showed individual repeatability over time (i.e. between 3 rounds over ∼6-8 week period). Similarly, all species, except for New Caledonian crows, showed individual repeatability across all 3 conditions. Regarding the lack of individual repeatability in New Caledonian crows, these were the only wild birds (temporarily captive) of the sample, which may have influenced their responses. It is also possible that this is related to habituation to the captive situation. Individual flexibility (i.e. lack of repeatability or inconsistency) may be more adaptive in the wild, where conditions can vary more widely than captivity. Additionally, individual inconsistency has been found in other corvid species, including pinyon jays and Clark’s nutcrackers exploratory responses to novel environments and novel objects (without familiar food present) ^16^. Some of these same individuals were tested in the present study, highlighting that neophobia may vary within and between individuals depending on types of neophobia, or aspects of study design, like task type.

The main limitations of this study, also applicable to some previous comparative cognition studies, were some unavoidable site differences. We therefore primarily used difference scores (novel condition minus control data) to aid in standardising latency scores across sites and control for baseline neophobia. We differentiated each of the socio-ecological factors tested on 2-levels, relying on published data to support these distinctions (e.g. ^27,38^), as it was not otherwise possible to determine each species reliably by other means. Some factors could be explored on further levels (such as a scale or distribution size for range) if supporting evidence becomes available for each species for such a distinction in future. There were differences in sample size per species, indicating care should be taken with any generalisations beyond the samples to wider species-levels. Our samples were also primarily captive individuals, which may influence neophobia ^43^. This study was a worthwhile and necessary first step into establishing a multi-lab collaboration, and captive birds allowed us to identify individuals, conduct repeated testing and control the environment, which could be expanded upon in future, for instance, to include corvids in the field ^7^. Being able to test more widely within groups of the same species from different backgrounds, as well as between species, and expanding these types of collaborative approaches to test other bird groups than corvids to explore the drivers of neophobia in birds more generally, is a recommended focus on future research. Furthermore, other aspects of neophobia, such as novel environments, predators or humans (e.g. ^25^) could be tested.

There are several wider implications of our study. When comparing neophobia in different species, it is important, where possible, to consider the role of socio-ecological factors, like diet, habitat use and sociality. Neophobia can influence how an animal interacts with novel problems, so should be tested as a baseline, particularly in new species/individuals, when conducting cognition research. The world is fast becoming more urbanised due to human activity, with many species being forced to adapt to changing environments or risk survival ^3^. As neophobia may impact how quickly a species or individual can adapt, it is a useful tool in designing conservation applications, such as in reintroductions ^21,34,44^. For example, the presentation of new bird feeders or safe nesting sites could be modified according to the species individual’s level of neophobia, and more neophobic individuals may require more pre-release training than others. Additionally, for species which are extinct in the wild, comparative behavioural and cognitive data from close relatives may help determine the extent to which long-term conservation breeding erodes natural responses. Therefore, neophobia and related research can provide valuable information in basic and applied research.

In conclusion, this study established a global collaborative network among corvid researchers to investigate the socio-ecological correlates of neophobia in these birds. Furthermore, neophobia can impact cognitive performance ^19,44^, but is often not tested or accounted for in comparative research – this study contributes to resolving this issue. It also contributes to a growing push to conduct multi-species comparisons while simultaneously facilitating other collaborative work between these labs in the future. Though species differences in neophobia are well-known among those working with corvids, they are more typically incorporated into study designs (for example, including a habituation phase to new stimuli) than studied in their own right or comparatively across different species. By investigating neophobia across species that vary in several socio-ecological factors and feature frequently in studies of behaviour and cognition, this study has broad implications for those interested in behavioural ecology, evolutionary biology, comparative psychology and other related fields.

## Acknowledgements

We thank the study funders: the European Research Council under the European Union’s Seventh Framework Programme (FP7/2007-2013)/ERC Grant Agreement No. 3399933 awarded to N.S.C., Career Support Fund (University of Cambridge) to R.M., Natural Science and Engineering Research Council Discovery grant (#4944-2017) and Canada Research Chair fund to D.M.K., DFG grant (number BR 5908/1-1) to K.F.B., JSPS (KAKENHI 16H06324, 20H01787) to E.I., 19J22654 to A.S., Keio University Grant-in-Aid for Innovative Collaborative Research Projects MKJ1905 to E.I., US National Science Foundation SES-1658837 to J.R.S., Austrian Science Fund (FWF) grants (W1262-B29, P33960-B) to T.B., and (P26806) to J.J.M.M., a Royal Society of New Zealand Rutherford Discovery Fellowship and a Prime Minister’s McDiarmid Emerging Scientist Prize to A.H.T., and National Natural Science Foundation of China (No. 31772470) to Z.L. The funders had no role in study design, data collection and analysis, decision to publish, or preparation of the manuscript. Thank you very much to Alizée Vernouillet and Camille Troisi for assisting in part of data collection with some of the Eurasian jays, to Camille Troisi for feedback on a manuscript draft, and to Ellen Skipper for assisting in video coding for some Eurasian jay data (N.S.C. Lab).

## Author contributions

R.M., M.LL., A.F. A.L.G., N.S.C. conceived the study idea and research design. R.M. and M.L.L. project managed the study. R.M., S.A.R. and J.R.S. analysed the data, R.M., M.L.L. and J.R.S produced the figures. R.M., A.F., K.F.B., E.G.P., K.G., L.B.L., A.L.G., Y.L., M.S., A.K., P.S., L.W., L.M.W., Y.Z. collected the data. R.M., A.F., I.C., E.G.P., A.L.G. coded the videos. R.M. and I.C. wrote the manuscript, with comments and feedback from all other authors. R.M., K.B., T.B., K.G., E.I., D.M.K., Z.L., A.N., J.R.S., A.H.T., N.S.C. provided funding to support the study.

## Declaration of interests

The authors declare no competing interests.

## STAR Methods

### Subjects

We tested 241 corvid subjects (141 males, 95 females, 5 unknown, primarily adult birds) across 10 species and 10 lab teams worldwide (S8 Table). The sample sizes ranged from 9 to 108 subjects per species (mean = 24; median = 15), depending on subject availability. All subjects could be identified individually (e.g. by coloured leg rings). Species tested were common ravens (n=15), carrion/ hooded crows (n=18), large-billed crows (n=13), New Caledonian crows (n=9), ‘Alala□ (n=108), Eurasian jays (n=24), pinyon jays (n=21), blue jays (n=9), Clark’s nutcrackers (n=10) and azure-winged magpies (n=14). Each lab housed their own species according to the ethical and housing conditions required within each country, with two labs holding more than 1 species, and 3 species each tested at two different sites (S8 Table). Individual labs were responsible for the data collection of their birds but were provided with the same protocols to ensure the methodology remained consistent and were in regular contact with the organising team.

These species differ in several specific socio-ecological factors (Table 1). Information was collated as to whether species occupied a broad or restricted range (e.g. island-living endemic species), use of urban habitats (as well as rural and suburban), whether they hunt live birds and mammals, live in territorial pairs (primarily throughout the year or seasonally) or within family groups (e.g. dominant breeding pair with offspring), average flock size (small = up to 100 individuals, large = over 100 individuals), whether they cache (hide food to return to later) large amounts of a specific food during certain seasons (specialised) or a variety of food across the year (moderate), and if they were from the Corvus genus or not ^27,38,45–47^.

### Apparatus/materials

There were three conditions: control (familiar food alone), novel food, and novel object (novel items beside familiar food). The familiar food (placed in a familiar food bowl) varied between bird groups, depending on the regular diet in each lab. The novel food consisted of jelly in 3cm^3^ blocks, also placed in a (different) familiar food bowl. There were three colours/flavours of jelly used: orange, purple/blackcurrant, and green/lemon & lime, which were presented individually across the three rounds. As the species typically have different diets, and the food needed to be equally novel for them all, a colourful, human-made food such as jelly provided an ideal option (with prior ethical approval including from a Home Office appointed Named Veterinary Surgeon, Cambridge University). The novel objects came in three variations, but all had the same properties: they were made of multiple items and textures, with no part that could look like eyes (to avoid resembling predators), and all contained the colours blue, yellow, green, and red ^34^. Part of the objects also had to be shiny, and the objects were all between one third and one half the size of the subject (so the size of the object itself varied with species; S3 Figure). All birds were tested in a feeding or testing compartment/cage, which varied in dimensions by lab, but gave the birds as much room as possible to avoid and/or approach stimuli. The testing area was familiar to the bird, or else the bird was habituated to the cage prior to testing.

### Procedure

The tests involved measuring behavioural responses to novel food and novel objects beside familiar food, in relation to baseline measures of familiar food only (control). Data collection took place outside of breeding season, with adult, captive individuals, other than the New Caledonian crows, which were wild birds temporarily held in captivity. For most species/groups, individuals were temporarily separated in visually isolated testing compartments, though typically not acoustically isolated i.e. could hear groupmates (‘Alala□ were left in their regularly housed social groups for tests to reduce stress, which were primarily 2-bird breeding pairs). Separation was achieved via voluntary participation in some labs (e.g. Eurasian jays, New Caledonian crows, common ravens, ‘Alala□, as well as – in T.B. & J.J.M.M. lab - carrion crows and azure-winged magpies), while the other birds were physically moved by an experimenter to the familiar testing area as per the typical testing procedures in each lab. The novel item (food or object) was placed beside the familiar food dish (20cm for larger species i.e. *Corvus* genus, 10cm for smaller species i.e. other species), with items placed in the same location (e.g. a table/ platform/ mesh wall – large enough so that the bird could approach slowly from more than a body length away) for all tests and individuals within each species. Where possible, the stimuli were present before the subject entered the testing compartment (all species except ravens). The test trial started when the subject entered the testing compartment (or experimenter left compartment). Each trial lasted a maximum of 10 minutes (600 seconds) or ended when the subject touched the familiar food (i.e. beak contacted food).

Each novel test ‘round’ was conducted 3 times with 1 trial per condition per round (i.e. 9 trials in total) to allow for testing for individual repeatability within and between conditions. The control trial was conducted within 48 hours of both novel tests, and all in the morning, without withholding of food before testing if possible. Each round of testing (1 trial each of food-control-object conditions) took place with approx. 2 weeks between each round i.e. week 1: food-control-object, week 3: food-control-object, week 5: food-control-object. Therefore, testing took approximately 6 weeks in total to complete per species/group. The order of presentation of the novel food and objects was counterbalanced across subjects, e.g. subject 1, round 1 – novel food type 1 (orange jelly), round 2 – type 2 (green jelly), round 3 – type 3 (purple jelly); subject 2, round 1 – type 3, round 2 – type 1, round 3 – type 2 etc. The testing schedule for half of the subjects was food-control-object in every round, and for the other half object-control-food in every round per group. All species were tested in all three conditions, except for the ‘Alala□s, which were tested in the familiar food and novel object conditions only ^34^ (due to Covid-19 pandemic limiting access for testing the novel food condition). Most individuals participated in all trials, with minimal missing data (S8 Table).

Our main measure was latency to touch familiar food signifying how long the individual took to touch a familiar, desirable food in the presence of a novel item. Any avoidance of the novel item (and thus familiar food) can then be interpreted as neophobia ^1^. Latency to touch familiar food was used (rather than latency to eat) to control for any potential doubt as to whether the bird swallowed the food.

### Data Analyses

Trials were recorded and all new videos (>1200 videos were newly collected; >650 ‘Alala□ videos were coded previously for ^34^ study) were coded in Solomon Coder. 12-15% of video trials for each species/group were coded by a second coder to ensure inter-rater reliability: ‘Alala□: intra-class correlation coefficient (ICC) = 0.956, CI =0.94-0.97, p < 0.001; all other species: ICC = 0.879, CI=0.804-0.925, p < 0.001). The full corresponding dataset for all analysis and the R script is available at: https://figshare.com/s/16a77c3ab4e7569f0d98

We had three main research questions and associated analyses: 1. species comparison 2. effect of socio-ecological factors 3. individual temporal and contextual repeatability of neophobia. The main dependent variable was latency to touch familiar food (0-600 seconds). We used R (version 4.1.0) for all analysis. For Q1: we conducted a Linear Mixed Model (LMM) to assess which factors influenced latency to touch familiar food. The residuals of a LMM visually approached normal distribution (although the Shapiro-Wilk test indicated the distribution was different from normal, W=0.9919, p<0.001). We compared the LMM (packages lm4, car, functions lmer(), anova(), and Anova()) with the raw latency scores with an LMM using a log (base 10) transformation of latency + 1 (to avoid 0s). A likelihood ratio test (using anova() function) showed that the log-transformed model was preferred over the raw latencies (AIC raw = 21934.6, AIC log10 = 2761.5). Further transformations and Generalized Linear Mixed Models with other error distributions and link functions did not improve model fit. We therefore used the log-transformed latencies for all analysis, though we plot the raw latencies for visual clarity. With all LMMs, we used likelihood ratio tests to investigate the effect of the individual predictors (using drop1() function with best-fit model as input and setting test statistic to chi-square). We used Tukey comparisons (package multcomp, function glht()) for post-hoc tests without direct p-value correction. P-value corrections, such as Bonferroni, limit the number of possible comparisons ^49^ and comparison of multiple species was a primary aim in this study.

In LMM 1, using all data, we included the main effects of condition, species, and round in the full model, with individual nested in site as a random effect and all variables set as factors. A potential confound of our study is that most species were housed and tested in differing locations and conditions, including testing compartment size. Site is therefore correlated closely with species. However, three species were tested at two locations; therefore, we checked these three species individually for an effect of site (LMM, site as main effects, individual as random effect; S2 Table).

To directly examine potential neophobia effects of novel objects and food, we calculated differences scores by subtracting the log-transformed latency values of the control condition from those of the novel object condition and separately for the novel food condition. Therefore, the control serves as the baseline for how long it usually takes an individual to touch familiar food (without novel items present). By subtracting this control value from the latency to touch familiar food when a novel object was present should help to standardize for any site differences like cage size, e.g. species A has a small test cage so may have a shorter control latency due to this (less space to cover/ more likely to be closer at the start of the test) compared with species B with a large test cage. We created pairwise individual difference scores for each round and individual (e.g. individual 1, novel object round 1 minus control round 1; novel object round 2 minus control round 2). In LMM 2 (object difference scores) and LMM 3 (food difference scores), we included the main effects of species and round, with individual nested in site as the random effect.

For Q2: we conducted LMM 4 (object difference score) and LMM 5 (food difference score), with the main effects of range, urban habitat, adult sociality, flock size, caching, live hunting, and genus, with individual nested in site as a random effect. The full models (including all predictor variables) had the best fit according to AIC. Though accounting for phylogenetic relationships can be important in some situations, testing for phylogenetic signal with fewer than 20 species is problematic ^41,42,50^, testing is not advisable for all research questions (e.g. Q1) ^50^, and the corvid evolutionary tree is not yet well established for all tested species (e.g. conflicting genetic results about the closest relative for ‘Alala□) ^51^. Therefore, we did not include a phylogenetic control in our analyses. We did, however, include the variable ‘genus’ (Corvus or not) in our Q2 models. Additionally, we provide a phylogenetic tree for visualisation purposes with relative neophobia scores per species (Figure 1). In reporting all results, we avoid using the term ‘significant’ ^52^.

For Q3, we tested across species and within species for individual repeatability over time (across rounds) and over context (across conditions) using intraclass correlation coefficients (ICCs). We extracted ICC estimates from linear models with individual as a random effect and bootstrapped 1,000 samples to generate 95% confidence intervals around the estimates (R package rpt, using rpt() function). For contextual repeatability, we included condition in the linear model, and for temporal repeatability, we included round in the model.

The ‘Alala□ control and novel object data was collected and examined in a previous study ^34^. We used a comparable methodology as this study while collecting all the new data with the 9 new corvid species for the present study. We edited the ‘Alala□ data set for the present study by introducing a cut-off of maximum of 10 minutes for each trial (original data set maximum of 60 min trials) – any individuals that did not touch familiar food within 10 minutes were assigned 600 seconds – to ensure comparability.

Example video trials can be found at: https://youtu.be/Lhzyk3srmdg.

### Ethics Statement

For animal research, all applicable international, national and/or institutional guidelines for the care and use of animals were followed. For N.S.C’s Comparative Cognition lab, this non-invasive behavioural study with birds was conducted adhering to UK laws and regulations and was covered under a non-regulated procedure through University of Cambridge, approved by the Home Office appointed Named Animal Care and Welfare Officer, Named Veterinary Surgeon and Chairperson for the Psychology and Zoology Department Animal User’s Management Committee. For D.M.K lab, research protocol approved by University of Manitoba’s Animal User Committee (F18-041) and complied with the guidelines set by the Canadian Council on Animal Care. For A.N., experiments were approved by the national authorities (Regierungspräsidium). For E.I. lab, the experimental protocol (number 9069) authorised by the Animal Care and Use Committee of Keio University, for capturing wild crows (numbers 27924005 and 29030001) authorised by the Japanese Ministry of the Environment. For J.R.S. lab, research protocol approved by University of Nebraska-Lincoln IACUC (number 1708). ForA.G. contribution, work was approved by San Diego Zoo Global’s animal care and use committee IACUC (number 16-009) and conducted under USFWS Permit (number TE-060179-5) and State of Hawaii Division of Forestry and Wildlife permit (number WL16-04). For K.G. lab, a research protocol approved by Luther College IACUC (no. 2019-4). For A.H.T. lab, a University of Auckland Animal Ethics Committee (no. 001823). For T.B. lab, work on foraging decisions, including this non-invasive behavioural study, was conducted adhering to Austrian law (2. Federal Law Gazette no. 501/1989) and approved by an Animal Ethics and Experimentation Board of the Faculty of Life Sciences, University of Vienna. For Z.L. lab, the study was conducted according to the Ethics Review Committee of Nanjing University (no. 2009-116), under Chinese law, no specific approval was required for this non-invasive study.

## Supplemental Information Legends

**S1 Figure. Latency to touch familiar food in each round, across all conditions and species**. Round 3 differs significantly from round 1 and 2, while round 1 and 2 do not differ significantly from each other. Points represent individuals, lines represent median. * p < 0.05

**S2 Figure. Site effect on latency to touch familiar food in azure-winged magpie, carrion crow and pinyon jay**

**S3 Figure. Example of novel objects for Eurasian jays**

**S1 Table. Pairwise comparisons of latency data between species**

**S2 Table. Linear mixed models with main effect of site on latency to touch familiar food for the three species that were tested in two sites**

**S3 Table. Pairwise comparisons of novel object difference scores between species**

**S4 Table. Pairwise comparisons of novel food difference scores between species**

**S5 Table. Individual temporal repeatability within each species and condition**

**S6 Table. Individual contextual repeatability within each species**

**S7 Table. Individual temporal and contextual repeatability**

**S8 Table. Subject information, including sex, source and participation in testing**

## Notes

### Competing Interest Statement

The authors have declared no competing interest.

### Summary of Updates

Correction to spelling of one author name and minor amendment to ethics permission for one lab.

