## Supplementary Materials for "Individual repeatability, species differences, and the influence of socio-ecological factors on neophobia in 10 corvid species" (from another version of this preprint: version 1, 2021-07-27)

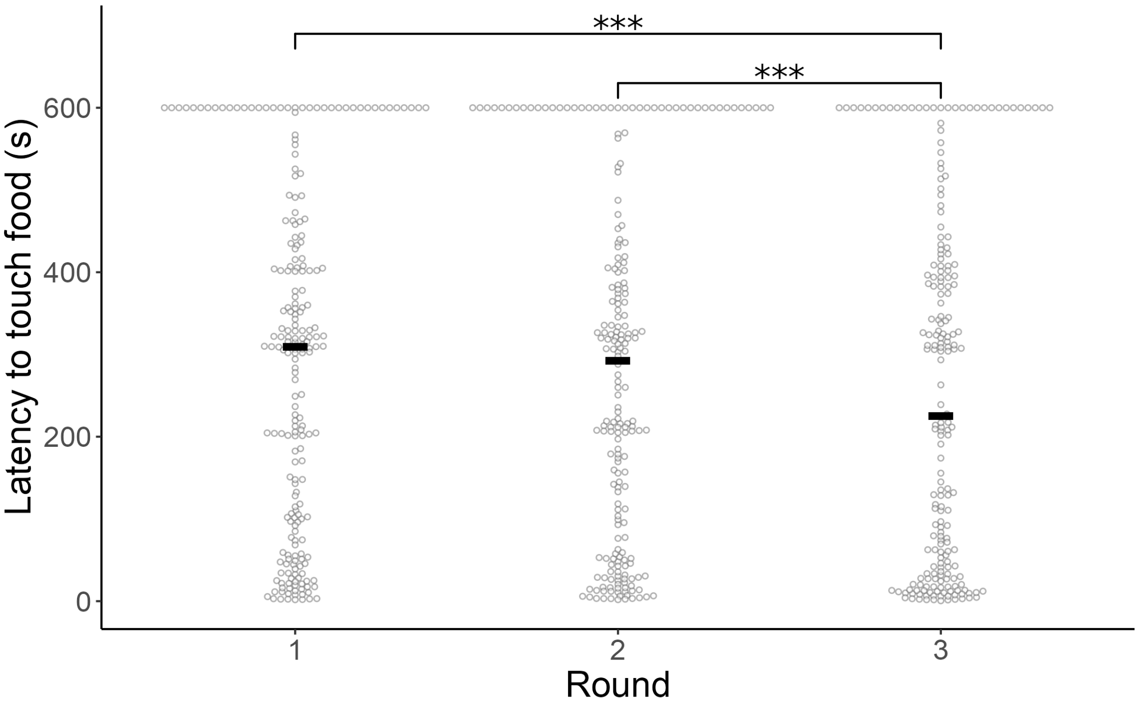


*Figure S1*. Latency to touch familiar food in each round, across all conditions and species. Round 3 differs from round 1 and 2, while round 1 and 2 do not differ from each other. Points represent individuals, lines represent median.

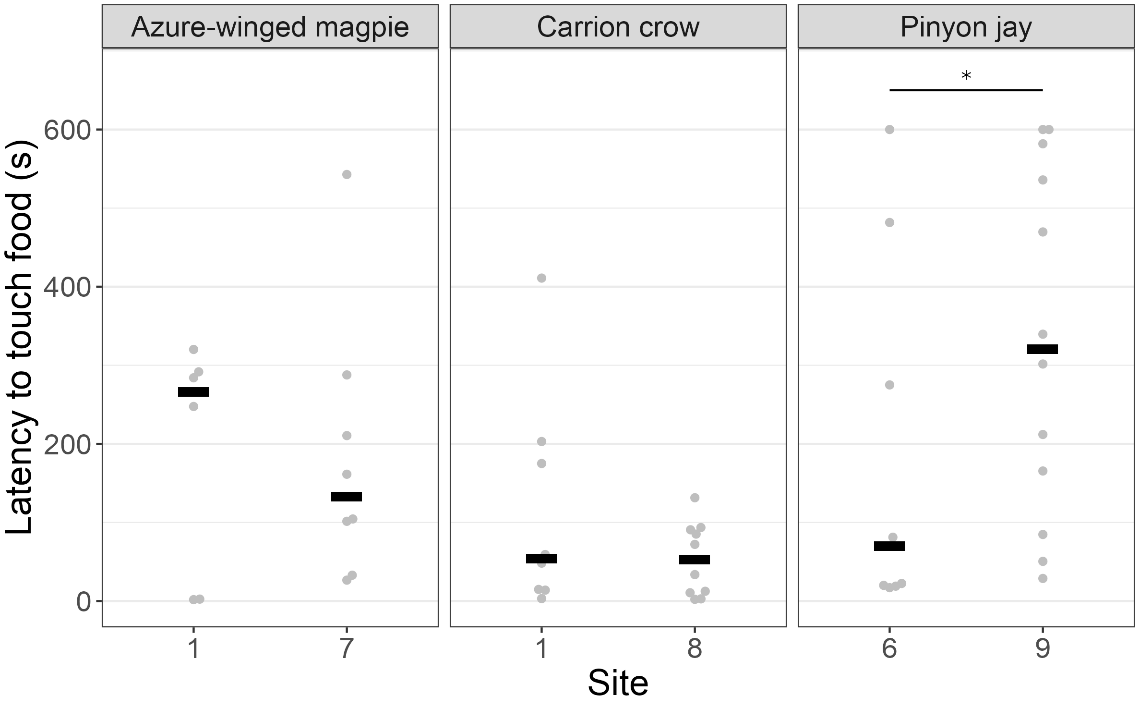


*Figure S2*. Site effect on latency to touch familiar food in azure-winged magpie, carrion crow and pinyon jay.

Table S1

*Pairwise comparisons of latency data between species*

| | Estimate | Standard error | z | p-value |
|---|---|---|---|---|
| Blue jay - Azure-winged magpie | 0.491 | 0.209 | 2.351 | 0.019 |
| Carrion crow - Azure-winged magpie | -0.496 | 0.177 | -2.811 | 0.005 |
| Clark's nutcracker - Azure-winged magpie | 0.518 | 0.203 | 2.558 | 0.011 |
| Common raven - Azure-winged magpie | -0.437 | 0.183 | -2.392 | 0.017 |
| Eurasian jay - Azure-winged magpie | 0.284 | 0.166 | 1.710 | 0.087 |
| 'Alalā - Azure-winged magpie | 0.416 | 0.144 | 2.891 | 0.004 |
| Large-billed crow - Azure-winged magpie | 0.668 | 0.189 | 3.540 | 0.000 |
| New Caledonian crow - Azure-winged magpie | -0.316 | 0.209 | -1.513 | 0.130 |
| Pinyon jay - Azure-winged magpie | 0.118 | 0.170 | 0.693 | 0.488 |
| Carrion crow - Blue jay | -0.988 | 0.199 | -4.959 | 0.000 |
| Clark's nutcracker - Blue jay | 0.027 | 0.223 | 0.122 | 0.903 |
| Common raven - Blue jay | -0.929 | 0.205 | -4.537 | 0.000 |
| Eurasian jay - Blue jay | -0.207 | 0.190 | -1.091 | 0.275 |
| 'Alalā - Blue jay | -0.076 | 0.171 | -0.443 | 0.658 |
| Large-billed crow - Blue jay | 0.177 | 0.210 | 0.843 | 0.399 |
| New Caledonian crow - Blue jay | -0.808 | 0.228 | -3.536 | 0.000 |
| Pinyon jay - Blue jay | -0.374 | 0.193 | -1.937 | 0.053 |
| Clark's nutcracker - Carrion crow | 1.015 | 0.193 | 5.271 | 0.000 |
| Common raven - Carrion crow | 0.059 | 0.172 | 0.344 | 0.731 |
| Eurasian jay - Carrion crow | 0.780 | 0.154 | 5.079 | 0.000 |
| 'Alalā - Carrion crow | 0.912 | 0.129 | 7.065 | 0.000 |
| Large-billed crow - Carrion crow | 1.165 | 0.178 | 6.547 | 0.000 |
| New Caledonian crow - Carrion crow | 0.180 | 0.199 | 0.904 | 0.366 |
| Pinyon jay - Carrion crow | 0.614 | 0.157 | 3.901 | 0.000 |
| Common raven - Clark's nutcracker | -0.956 | 0.198 | -4.822 | 0.000 |
| Eurasian jay - Clark's nutcracker | -0.234 | 0.183 | -1.281 | 0.200 |
| 'Alalā - Clark's nutcracker | -0.103 | 0.163 | -0.631 | 0.528 |
| Large-billed crow - Clark's nutcracker | 0.150 | 0.204 | 0.736 | 0.462 |
| New Caledonian crow - Clark's nutcracker | -0.835 | 0.223 | -3.749 | 0.000 |
| Pinyon jay - Clark's nutcracker | -0.401 | 0.186 | -2.154 | 0.031 |
| Eurasian jay - Common raven | 0.721 | 0.161 | 4.487 | 0.000 |
| 'Alalā - Common raven | 0.853 | 0.137 | 6.206 | 0.000 |
| Large-billed crow - Common raven | 1.106 | 0.184 | 6.007 | 0.000 |
| New Caledonian crow - Common raven | 0.121 | 0.205 | 0.591 | 0.554 |
| Pinyon jay - Common raven | 0.555 | 0.164 | 3.377 | 0.001 |
| 'Alalā - Eurasian jay | 0.132 | 0.114 | 1.151 | 0.250 |
| Large-billed crow - Eurasian jay | 0.384 | 0.168 | 2.294 | 0.022 |
| New Caledonian crow - Eurasian jay | -0.600 | 0.190 | -3.159 | 0.002 |
| Pinyon jay - Eurasian jay | -0.166 | 0.146 | -1.144 | 0.253 |
| Large-billed crow - 'Alalā | 0.253 | 0.145 | 1.739 | 0.082 |
| New Caledonian crow - 'Alalā | -0.732 | 0.171 | -4.286 | 0.000 |
| Pinyon jay - 'Alalā | -0.298 | 0.119 | -2.499 | 0.012 |
| New Caledonian crow - Large-billed crow | -0.985 | 0.210 | -4.686 | 0.000 |
| Pinyon jay - Large-billed crow | -0.551 | 0.171 | -3.222 | 0.001 |
| Pinyon jay - New Caledonian crow | 0.434 | 0.193 | 2.247 | 0.025 |

Table S2

*Linear mixed models with main effect of site on latency to touch familiar food for the three species that were tested in two sites*

| | Azure-winged magpie | | | Carrion crow | | | Pinyon jay | | |
|---|---|---|---|---|---|---|---|---|---|
| Fixed effect | Chisq | df | p-value | Chisq | df | p-value | Chisq | df | p-value |
| site | 1.024 | 1 | 0.312 | 0.078 | 1 | 0.78 | 4.35 | 1 | 0.037 |

Table S3

*Pairwise comparisons of novel object difference scores between species*

| | Estimate | Standard error | z | p-value |
|---|---|---|---|---|
| Blue jay - Azure-winged magpie | -0.183 | 0.202 | -0.905 | 0.366 |
| Carrion crow - Azure-winged magpie | 0.193 | 0.175 | 1.101 | 0.271 |
| Clark’s nutcracker - Azure-winged magpie | -0.307 | 0.196 | -1.566 | 0.117 |
| Common raven - Azure-winged magpie | 0.396 | 0.178 | 2.227 | 0.026 |
| Eurasian jay - Azure-winged magpie | 0.191 | 0.164 | 1.165 | 0.244 |
| ’Alalā - Azure-winged magpie | 0.248 | 0.140 | 1.777 | 0.076 |
| Large-billed crow - Azure-winged magpie | -0.438 | 0.183 | -2.397 | 0.017 |
| New Caledonian crow - Azure-winged magpie | -0.134 | 0.203 | -0.658 | 0.510 |
| Pinyon jay - Azure-winged magpie | -0.059 | 0.165 | -0.358 | 0.720 |
| Carrion crow - Blue jay | 0.375 | 0.193 | 1.942 | 0.052 |
| Clark’s nutcracker - Blue jay | -0.124 | 0.212 | -0.584 | 0.559 |
| Common raven - Blue jay | 0.579 | 0.196 | 2.955 | 0.003 |
| Eurasian jay - Blue jay | 0.373 | 0.183 | 2.040 | 0.041 |
| ’Alalā - Blue jay | 0.431 | 0.162 | 2.660 | 0.008 |
| Large-billed crow - Blue jay | -0.255 | 0.200 | -1.275 | 0.202 |
| New Caledonian crow - Blue jay | 0.049 | 0.219 | 0.224 | 0.823 |
| Pinyon jay - Blue jay | 0.124 | 0.184 | 0.671 | 0.502 |
| Clark’s nutcracker - Carrion crow | -0.499 | 0.187 | -2.670 | 0.008 |
| Common raven - Carrion crow | 0.204 | 0.168 | 1.210 | 0.226 |
| Eurasian jay - Carrion crow | -0.002 | 0.153 | -0.014 | 0.989 |
| ’Alalā - Carrion crow | 0.056 | 0.127 | 0.438 | 0.661 |
| Large-billed crow - Carrion crow | -0.631 | 0.173 | -3.639 | 0.000 |
| New Caledonian crow - Carrion crow | -0.326 | 0.194 | -1.678 | 0.093 |
| Pinyon jay - Carrion crow | -0.252 | 0.154 | -1.632 | 0.103 |
| Common raven - Clark’s nutcracker | 0.703 | 0.190 | 3.704 | 0.000 |
| Eurasian jay - Clark’s nutcracker | 0.497 | 0.176 | 2.819 | 0.005 |
| ’Alalā - Clark’s nutcracker | 0.555 | 0.155 | 3.590 | 0.000 |
| Large-billed crow - Clark’s nutcracker | -0.132 | 0.194 | -0.677 | 0.498 |
| New Caledonian crow - Clark’s nutcracker | 0.173 | 0.213 | 0.811 | 0.417 |
| Pinyon jay - Clark’s nutcracker | 0.248 | 0.178 | 1.394 | 0.163 |
| Eurasian jay - Common raven | -0.206 | 0.156 | -1.317 | 0.188 |
| ’Alalā - Common raven | -0.148 | 0.131 | -1.127 | 0.260 |
| Large-billed crow - Common raven | -0.834 | 0.176 | -4.733 | 0.000 |
| New Caledonian crow - Common raven | -0.530 | 0.197 | -2.689 | 0.007 |
| Pinyon jay - Common raven | -0.455 | 0.158 | -2.890 | 0.004 |
| ’Alalā - Eurasian jay | 0.058 | 0.111 | 0.522 | 0.602 |
| Large-billed crow - Eurasian jay | -0.629 | 0.162 | -3.886 | 0.000 |
| New Caledonian crow - Eurasian jay | -0.324 | 0.184 | -1.760 | 0.078 |
| Pinyon jay - Eurasian jay | -0.250 | 0.141 | -1.768 | 0.077 |
| Large-billed crow - ’Alalā | -0.687 | 0.138 | -4.985 | 0.000 |
| New Caledonian crow - ’Alalā | -0.382 | 0.163 | -2.338 | 0.019 |
| Pinyon jay - ’Alalā | -0.307 | 0.113 | -2.727 | 0.006 |
| New Caledonian crow - Large-billed crow | 0.304 | 0.201 | 1.512 | 0.131 |
| Pinyon jay - Large-billed crow | 0.379 | 0.163 | 2.325 | 0.020 |
| Pinyon jay - New Caledonian crow | 0.075 | 0.185 | 0.402 | 0.687 |

Table S4

*Pairwise comparisons of novel food difference scores between species*

| | Estimate | Standard error | z | p-value |
|---|---|---|---|---|
| Blue jay - Azure-winged magpie | -0.005 | 0.179 | -0.031 | 0.976 |
| Carrion crow - Azure-winged magpie | 0.145 | 0.156 | 0.932 | 0.352 |
| Clark's nutcracker - Azure-winged magpie | -0.010 | 0.173 | -0.059 | 0.953 |
| Common raven - Azure-winged magpie | 0.010 | 0.158 | 0.066 | 0.948 |
| Eurasian jay - Azure-winged magpie | 0.446 | 0.146 | 3.046 | 0.002 |
| Large-billed crow - Azure-winged magpie | 0.012 | 0.162 | 0.072 | 0.942 |
| New Caledonian crow - Azure-winged magpie | -0.186 | 0.179 | -1.040 | 0.298 |
| Pinyon jay - Azure-winged magpie | 0.024 | 0.146 | 0.164 | 0.869 |
| Carrion crow - Blue jay | 0.151 | 0.171 | 0.879 | 0.380 |
| Clark's nutcracker - Blue jay | -0.005 | 0.187 | -0.025 | 0.980 |
| Common raven - Blue jay | 0.016 | 0.173 | 0.091 | 0.927 |
| Eurasian jay - Blue jay | 0.452 | 0.163 | 2.771 | 0.006 |
| Large-billed crow - Blue jay | 0.017 | 0.177 | 0.097 | 0.923 |
| New Caledonian crow - Blue jay | -0.180 | 0.192 | -0.937 | 0.349 |
| Pinyon jay - Blue jay | 0.029 | 0.163 | 0.181 | 0.856 |
| Clark's nutcracker - Carrion crow | -0.155 | 0.166 | -0.936 | 0.349 |
| Common raven - Carrion crow | -0.135 | 0.150 | -0.900 | 0.368 |
| Eurasian jay - Carrion crow | 0.301 | 0.138 | 2.187 | 0.029 |
| Large-billed crow - Carrion crow | -0.133 | 0.154 | -0.867 | 0.386 |
| New Caledonian crow - Carrion crow | -0.331 | 0.171 | -1.930 | 0.054 |
| Pinyon jay - Carrion crow | -0.121 | 0.137 | -0.883 | 0.377 |
| Common raven - Clark's nutcracker | 0.021 | 0.168 | 0.122 | 0.903 |
| Eurasian jay - Clark's nutcracker | 0.456 | 0.157 | 2.902 | 0.004 |
| Large-billed crow - Clark's nutcracker | 0.022 | 0.172 | 0.127 | 0.899 |
| New Caledonian crow - Clark's nutcracker | -0.175 | 0.187 | -0.936 | 0.349 |
| Pinyon jay - Clark's nutcracker | 0.034 | 0.157 | 0.218 | 0.827 |
| Eurasian jay - Common raven | 0.436 | 0.140 | 3.113 | 0.002 |
| Large-billed crow - Common raven | 0.001 | 0.156 | 0.009 | 0.993 |
| New Caledonian crow - Common raven | -0.196 | 0.173 | -1.131 | 0.258 |
| Pinyon jay - Common raven | 0.014 | 0.139 | 0.098 | 0.922 |
| Large-billed crow - Eurasian jay | -0.435 | 0.145 | -3.007 | 0.003 |
| New Caledonian crow - Eurasian jay | -0.632 | 0.163 | -3.876 | 0.000 |
| Pinyon jay - Eurasian jay | -0.422 | 0.127 | -3.337 | 0.001 |
| New Caledonian crow - Large-billed crow | -0.197 | 0.177 | -1.116 | 0.265 |
| Pinyon jay - Large-billed crow | 0.012 | 0.144 | 0.085 | 0.932 |
| Pinyon jay - New Caledonian crow | 0.210 | 0.163 | 1.290 | 0.197 |

Table S5

*Individual temporal repeatability within each species and condition*

| Species | Control | Novel object | Novel food |
|---|---|---|---|
| Azure-winged magpie | N=13, ICC=0.309 (0.000, 0.658), p = 0.044 | N=13, ICC=0.593 (0.211, 0.827), p = 0.001 | N=14, ICC=0.233 (0.000, 0.611), p = 0.121 |
| Blue jay | N=9, ICC=0.962 (0.864, 0.989), p < 0.001 | N=9, ICC=0.814 (0.432, 0.939), p < 0.001 | N=9, ICC=0.680 (0.252, 0.898), p < 0.001 |
| Carrion crow | N=18, ICC=0.740 (0.477, 0.884), p < 0.001 | N=17, ICC=0.755 (0.536, 0.893), p < 0.001 | N=17, ICC=0.456 (0.102, 0.731), p = 0.002 |
| Clark's nutcracker | N=10, ICC=0.535 (0.100, 0.818), p = 0.002 | N=10, ICC=0.582 (0.134, 0.832), p = 0.001 | N=10, ICC=0.492 (0.043, 0.801), p = 0.005 |
| Common raven | N=15, ICC=0.638 (0.306, 0.835), p < 0.001 | N=15, ICC=0.707 (0.436, 0.878), p < 0.001 | N=15, ICC=0.630 (0.302, 0.828), p < 0.001 |
| Eurasian jay | N=24, ICC=0.557 (0.289, 0.754), p < 0.001 | N=24, ICC=0.290 (0.000, 0.568), p = 0.012 | N=24, ICC=0.346 (0.039, 0.634), p = 0.003 |
| 'Alalā | N=84, ICC=0.542 (0.418, 0.650), p < 0.001 | N=84, ICC=0.540 (0.426, 0.653), p < 0.001 | NA |
| Large-billed crow | N=13, ICC=0.496 (0.117, 0.761), p = 0.001 | N=13, ICC=0.216 (0.000, 0.571), p = 0.099 | N=13, ICC=0.426 (0.048, 0.734), p = 0.006 |
| New Caledonian crow | N=9, ICC=0.177 (0.000, 0.610), p = 0.189 | N=9, ICC=0.164 (0.000, 0.625), p = 0.245 | N=9, ICC=0.000 (0.000, 0.421), p = 1.000 |
| Pinyon jay | N=21, ICC=0.690 (0.464, 0.842), p < 0.001 | N=21, ICC=0.863 (0.733, 0.934), p < 0.001 | N=21, ICC=0.738 (0.535, 0.861), p < 0.001 |

Table S6

*Individual contextual repeatability within each species*

| Species | Repeatability across conditions |
|---|---|
| Azure-winged magpie | N=14, ICC=0.740 (0.438, 0.892), p < 0.001 |
| Blue jay | N=9, ICC=0.889 (0.646, 0.966), p < 0.001 |
| Carrion crow | N=18, ICC=0.721 (0.486, 0.873), p < 0.001 |
| Clark's nutcracker | N=10, ICC=0.693 (0.292, 0.891), p < 0.001 |
| Common raven | N=15, ICC=0.802 (0.588, 0.910), p < 0.001 |
| Eurasian jay | N=24, ICC=0.583 (0.330, 0.763), p < 0.001 |
| 'Alalā | N=84, ICC=0.545 (0.384, 0.679), p < 0.001 |
| Large-billed crow | N=13, ICC=0.651 (0.328, 0.849), p < 0.001 |
| New Caledonian crow | N=9, ICC=0.270 (0.000, 0.687), p = 0.091 |
| Pinyon jay | N=21, ICC=0.800 (0.640, 0.905), p < 0.001 |

Table S7

*Individual temporal and contextual repeatability*

| Control | Novel food | Novel object | Overall repeatability |
|---|---|---|---|
| N=216, ICC=0.542 (0.467, 0.625), p < 0.001 | N=132, ICC=0.477 (0.381, 0.591), p < 0.001 | N=215, ICC=0.548 (0.469, 0.625), p < 0.001 | N=217, ICC=0.462 (0.402, 0.521), p < 0.001 |

Table S8

*Subject information, including sex, source and participation in testing*

| Species | Sex (male.female or unknown) | Rearing information | Housing | Completed testing | Lab leader and location |
|---|---|---|---|---|---|
| Common raven, *Corvus corax* | 7.8 | Hand-reared | Group/ pair | 1 missing trial for round 2 & 3 for all conditions (did not participate) | Thomas Bugnyar, Austria |
| Carrion crow, *C. corone; C. cornix* | 4.4 | Hand-reared | Group/ pair | 4 missing trials for control round 2 & 3 and food round 3 and object round 3; 1 missing trial from food round 1; 2 missing trials from food round 2 and object round 2 (did not participate) | Thomas Bugnyar, Austria |
| Carrion crow, *C. corone* | 10.0 | Hand-reared | Group | Yes | Andreas Nieder, Germany |
| Large-billed crow, *C. macrorhynchos* | 8.5 | Wild: parent reared | Group | Yes | Ei-Ichi Izawa, Japan |
| New Caledonian crow, *C. moneduloides* | 3.6 | Wild: parent reared | Group | Yes | Alexander Taylor, New Zealand (subjects in New Caledonia) |
| ’Alalā, *C. hawaiiensis* | 64.44 | Hand-reared and parent-reared | Group/ pair | Control and novel object conditions only (not food condition) | Alison Greggor, USA |
| Eurasian jay, *Garrulus glandarius* | 14.10 | Hand-reared | Group | 2 missing trials for control round 2; 5 missing trails for control round 3; 4 missing trials for food round 2 & 3 and object round 2; with 2 missing trials in object round 3 (did not participate) | Nicola Clayton, UK |
| Pinyon jay, *Gymnorhinus cyanocephalus* | 10.2 | Wild: parent reared | Individual cages | Yes | Jeffrey Stevens, USA |
| Pinyon jay, *Gymnorhinus cyanocephalus* | 4.5 | Wild: parent reared | Individual cages | Yes | Debbie Kelly, Canada |
| Blue jay, *Cyanocitta cristata* | 2.2 (5 unknown) | Hand-reared | Individual cages | Yes | Kristy Gould, USA |
| Clark’s nutcracker, *Nucifraga columbiana* | 7.3 | Wild: parent reared | Individual cages | Yes | Debbie Kelly, Canada |
| Azure-winged magpie, *Cyanopica cyanus* | 6.2 | Parent-reared & Hand-reared | Group | Yes | Zhongqiu Li, China |
| Azure-winged magpie, *Cyanopica cyanus* | 2.4 | Parent-reared & Hand-reared | Group | 2 missing trials for round 2 and 3 for all rounds (2 birds died due to unrelated causes) | Thomas Bugnyar & Jorg J.M. Massen, Austria |
